# Lithography-free Water Stable Conductive Polymer Nanowires

**DOI:** 10.1101/2025.01.08.631660

**Authors:** Damien Hughes, Abdelrazek H. Mousa, Chiara Musumeci, Malte Larsson, Muhammad Anwar Shameem, Umut Aydemir, Ludwig Schmiderer, Jonas Larsson, Magnus Berggren, Fredrik Ek, Roger Olsson, Martin Hjort

## Abstract

Free-standing nanowires can gain intracellular access without causing cellular stress or apoptosis. Current approaches to generate nanowires focus on lithographic patterning and inorganic materials (Si, GaAs, Al2O3, etc.) while organic materials are less explored. Use of organic conductive polymers allows for creation of soft mixed ion–electron conducting nanowires. Processing conductive polymers into nanowires is challenging due to the harsh chemicals and processing conditions used. Here, we demonstrate a lithography-free and scalable method to generate all-organic water-stable nanowires composed of conductive polymers. A nanoporous membrane is filled with conductive polymer in solution followed by a cross-linking step to make the polymer water stable. The surface of the membrane is anisotropically etched using a reactive ion etcher to reveal the polymer inside the pores, which extend from the membrane as nanowires. We interface the nanowires with model algal cells and human primary hematopoietic stem and progenitor cells.

**TOC Graphic:** 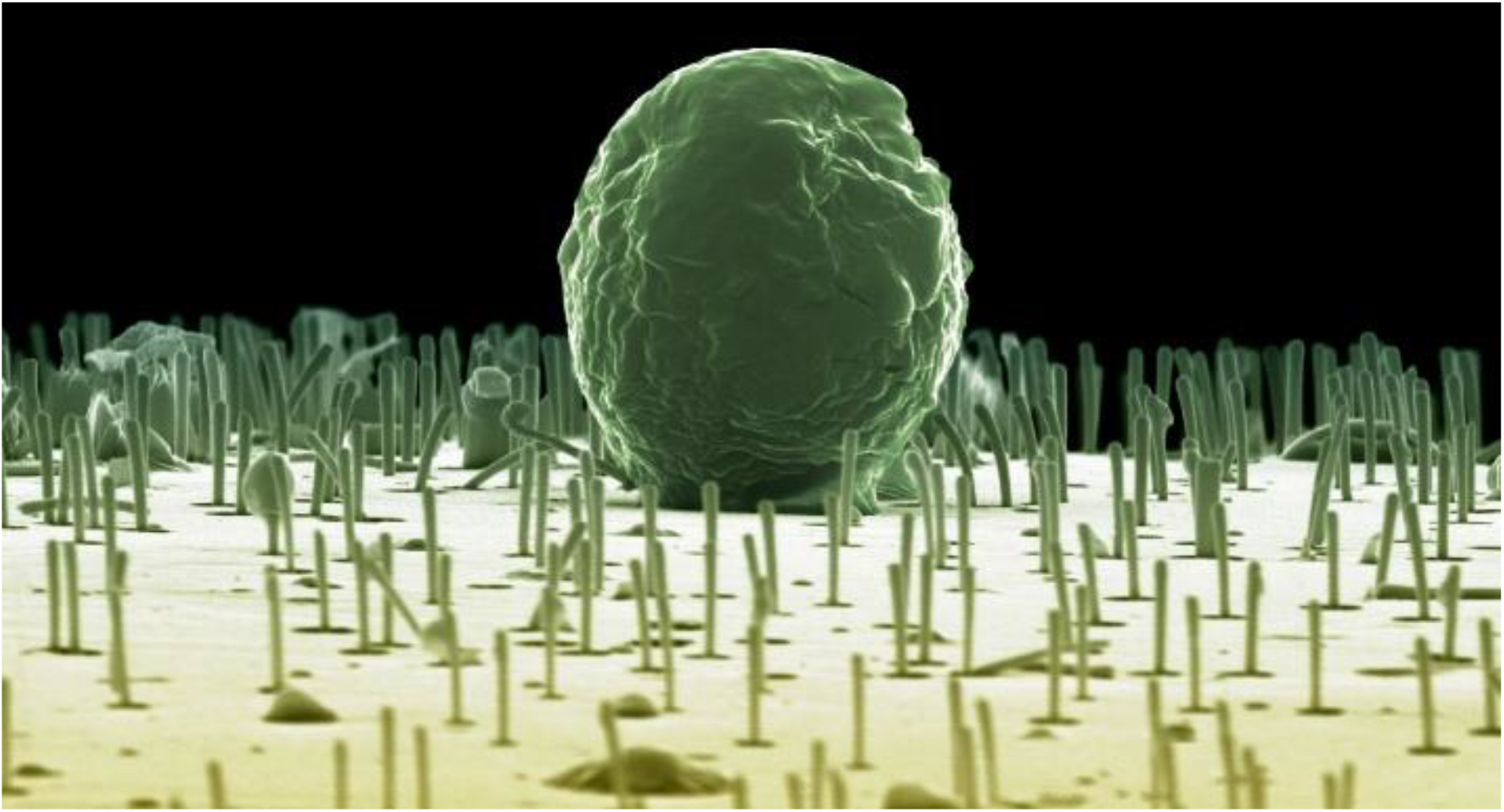

## Introduction

Free-standing, axially elongated nanostructures, such as nanowires (NWs), are ideal for applications within solar cells^1^, light-emitting diodes^2^, and electronics^3^. Lately, such nanostructures have been interfaced with living cells, making it possible to measure cellular traction forces^4^, deliver biomolecules into or out of living cells without toxic effects^5, 6^, and for artificial photosynthesis^7^. Due to their comparable ease of processability, these applications have so far been centered on nanostructures based on semiconductors^8, 9^, metals^10, 11^, or metal-oxides^12, 13^. A biocompatible carbon-based alternative is lacking.

Conductive polymers (CPs) are a class of materials being pursued for applications within bioelectronics due to their ability for mixed ion–electron conductivity^14, 15^ and their soft, self-healable, flexible nature which allows for seamless interfacing with tissue and cell cultures^4^. To comply with future needs, and to make use of the very high surface-to-volume ratio provided by nanostructure geometry, miniaturization is needed while still being able to pattern large areas.

Conventional photo- and electron lithography is poorly suited for this task due to harsh etching conditions and a lack of possibility to clean the CP from photoresist and other contaminants^16^.

Here, we present a lithography-free method to make a bespoke CP NW platform which can be used to interface with living cells. The NWs are formed by poly(3,4-ethylenedioxythiophene)butoxy-1-sulfonate (PEDOT-S)^17^ templated in track-etched (TE) membranes and made water-stable through introduction of cations or by electrofunctionalization with small thiophene trimers^18^. The methods allow full control over geometry and chemistry of the NWs by choosing different templates, processing parameters, and incorporation of trimers with PEDOT-S. The NW dimensions and densities are ideal for interfacing with cells^19, 20^ and even potential intracellular access.

## Results

### NW Processing

The NW processing starts with a commercially available TE polyimide (PI) or polycarbonate (PC) membrane (It4IP) serving as a template, Figure 1A. The TE membrane is coated with CP followed by a gas-phase etch step, which selectively removes the template membrane leaving the CP inside the pores to protrude as NWs. All dimensions of the NWs can be controlled by choosing a suitable starting membrane (density and diameter) and etch parameters (length).

**Figure 1:**
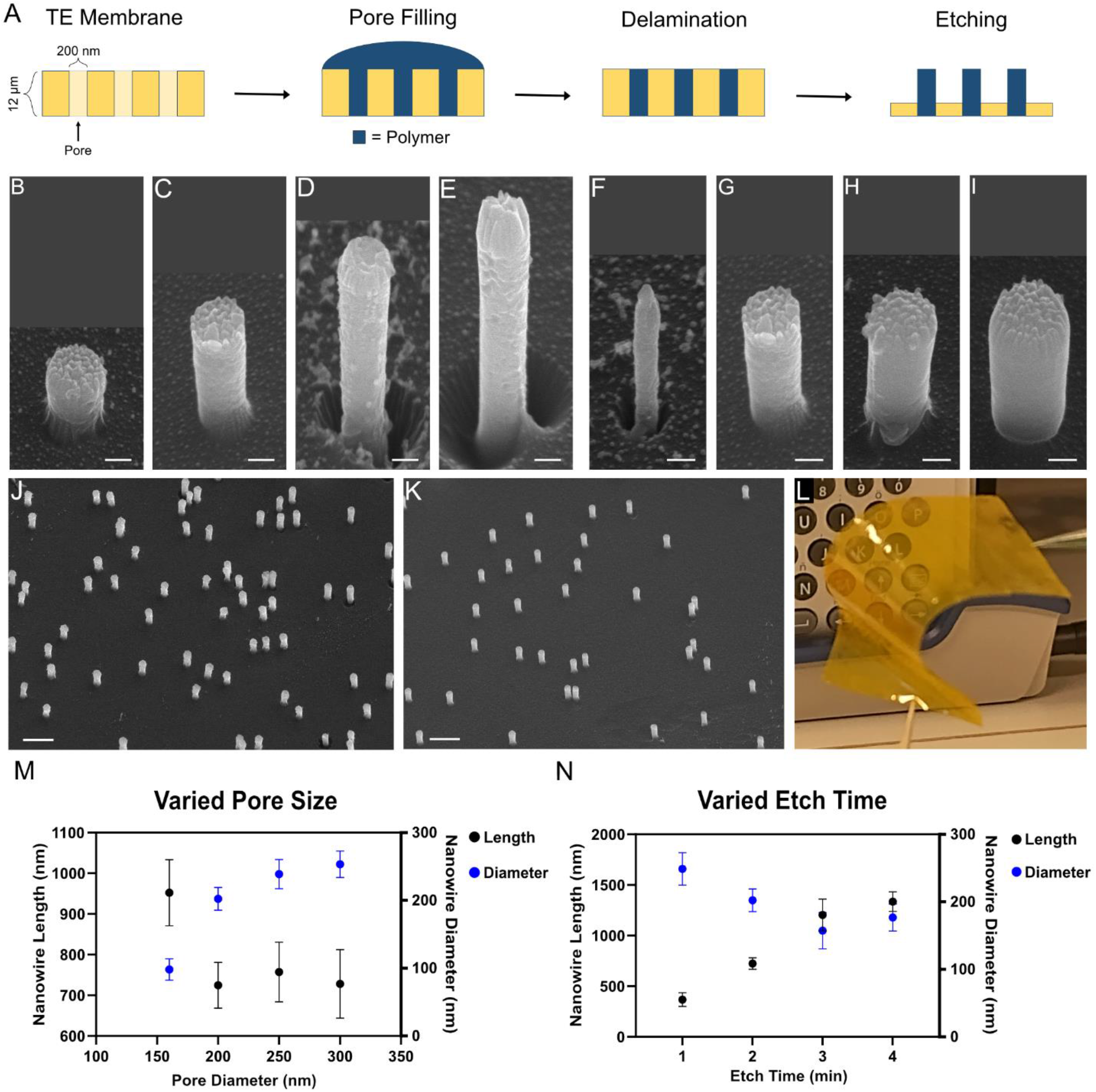
NW processing. (A) Schematic side-view describing the processing. A track-etched (TE) membrane is filled with CP solution and then allowed to dry. After drying, excess CP is removed. The TE membrane is then dry-etched using oxygen-based inductively coupled plasma reactive ion etching (ICP-RIE) to reveal the NWs. Please note that schematic is not to scale. (B– E) 30° tilted view SEM images of NWs of various lengths. Length can be controlled by modifying the etch time. Etching times used were 1 minute (B), 2 minutes (C), 3 minutes (D), and 4 minutes (E). (F–I) 30° tilted view SEM images of NWs made to have different thicknesses due to varying pore diameter. The pore diameters used were 160 nm (F), 200 nm (G), 250 nm (H), and 300 nm (I). (J–K) 30° tilted SEM images showing different pore densities, 5.5 e^7^ cm^−2^ for panel J and 2 e^7^ cm^−2^ for panel K. (L) Photograph of processed NWs on polyimide membrane. The photograph showcases both flexibility and transparency of the material. (M–N) Graphs illustrating the effects of pore size (M) and etch time (N) on NW length and diameter. The dots on the graph represent the mean (14–33 NWs measured per dot), while the error bars represent the standard deviation. The scale bars are 100 nm in B–I and 1 µm in J and K.

Membranes suitable for cellular interfacing have pore densities in the 10^7^ cm^−2^ range and thicknesses of at least 10 µm to ensure mechanical stability. The thickness of the membrane produces samples with high flexibility and optical transparency making it possible to image through the membrane, Figure 1L. While the pore densities can determine how cells interface with the NWs. Higher pore densities result in denser NWs and cells cultured on top of them will experience a bed-of-nails effect^21^ prohibiting a tight cell–NW interface whereas lower densities risk some cells not being able to reach a NW. Since non-carbon NWs with a diameter in the range of 100–300 nm have been found to establish healthy connections to cells^22, 6^ the same diameter range was chosen in this project.

Once the optimal TE membrane is determined, CPs in solution are added to the TE membrane to fill the pores. Pores become completely filled due to capillary forces. The physicochemical properties of the polymer solution are critical for efficient pore filling, guiding us to the use of the self-doped PEDOT-S, which contains a covalently attached sulfonate group for charge injection. The PEDOT-S must be well-dispersed and highly water soluble so as not to clog the pores or the pore openings. The most commonly used CP, poly(3,4-ethylenedioxythiophene):polystyrene-sulfate (PEDOT:PSS), did not provide a sufficient pore filling. This is presumably due to the PSS which is comparably larger than PEDOT, and can act to clog the pores. Further, even if PEDOT:PSS could fill the pores, there is a significant risk of phase separation where the PSS contents at different heights of the pores would vary (and thus alter the conductive properties).

Previously, we have used a high concentration of PEDOT-S (20 mg/ml) to install substrate-free electrodes in animal models^18, 23^. While that concentration was possible to inject using very thin capillaries (30 µm), it could not efficiently fill the nanopores (100–300 nm) for the NW templating used here. To conserve reagents and achieve less aggregation in the CP solution we used PEDOT-S at 5–10 mg/ml which resulted in NWs, Figure S1.

Once the polymer solution is applied, the sample is dried at room temperature before further processing. This leaves dried CP in the pores and on the top of the sample. The dry top layer is mechanically delaminated using masking tape, Figure 1A, resulting in CP only being located within the pores. Lastly, the membrane is brought into an inductively coupled plasma reactive ion etcher (ICP-RIE), a dry-etcher commonly used within semiconductor processing. In ICP-RIE, a plasma is generated in vacuum and high-energy ions impinge on the sample to react with it. The CP inside the pores etch at a lower rate than the membrane resulting in an isotropic etch where the NWs are seen to extend from the membrane. While this is the basic procedure for preparing NWs, modifications can be made to fit NWs to a specific need.

The length of the NWs can be tuned by using longer etch times or by increasing the ICP-RIE power input. We found that 25 W RF power and 500 W ICP was high enough to have a useful etch rate while maintaining membrane integrity. It is important to minimize the thermal load on the sample to avoid wrinkling or degradation of the plastic membrane. This is especially important for polycarbonate (glass transition temperature, Tg, 100–150 °C) but not as critical for polyimide (Tg > 300 °C). Figure 1B–E depicts representative NWs etched for 1–4 min showing a linear increase in length over time (320 nm/min). During longer etches, the NW diameter was observed to decrease (20 nm/min) allowing us to deduce a high etch selectivity of 16. Very long NWs reaching over 4 µm in length could also be processed without compromising NW structure (Figure S2).

NW diameter and density can be modified by choosing template membranes with different pore geometries. Nominal pore diameters, as defined by the vendor, was varied in between 160 and 300 nm (Figure 1F–I) and resulted in NWs ranging between 100 and 250 nm. The discrepancy stems from the vendors defining the average pore diameter in the membrane, whereas the NWs will only display the diameter at the top of the membrane, not taking conicity of the pores into account. NWs made using the 160 nm pore membrane were slightly longer than ones made from the other membranes indicating that porosity may affect the etch rate. Lastly, the density of NWs could be increased by using membranes with varying pore densities, Figure 1J–K, making them suitable to interface with cells of different sizes.

### Water Stability

To interface with cells or biological fluids the NWs must be water stable. While the NWs remain stable in organic solvents (e.g., acetone, isopropanol, ethanol, etc.), they readily dissolve in aqueous environments due to the high water-solubility of the PEDOT-S. To combat this, we have devised two routes to increase the water stability: addition of cations or electrofunctionalization using thiophene trimers.

Metal cations increase the water stability of PEDOT-S through ionic cross linking between the metal cation and the sulfonate group of PEDOT-S^17^ where higher valency cations exert more potent cross linking. We added cations to the CP solution before placing it onto the porous membrane. Trivalent cations, such as Fe^3+^, are the most efficient at stabilizing and can be added at mM concentrations without significantly affecting NW processing, Figure S1. The addition of trivalent cations is sufficient to increase water stability of NWs for up to 24 hours when placed into PBS (Figure S4). Divalent cations such as Ca^2+^, Cu^2+^, and Fe^2+^ could also be used to generate NWs, Figure S3. However, care must be taken to balance solubility, aggregation, and viscosity. Too high of an ion concentration combined with high PEDOT-S concentrations leads to polymer aggregating in the solution before addition to the TE membrane, leading to deficient pore filling (and thus no NWs), whereas too low ion concentration results in NWs which dissolve in water post-processing (Figure S4).

Another route to increase water stability as well as to modulate the volume properties of the NWs is by incorporating small, non-toxic thiophene-based trimers^23, 18, 24, 25^. The trimers are composed of two ethylenedioxythiophenes (EDOTs) with a functionalized thiophene in the middle. The trimer used here, 4-(2-(2,5-bis(2,3-dihydrothieno[3,4-b][1,4]dioxin-5-yl)thiophene-3-yl)ethoxy)butane-1-sulfonate (ETE-S) is functionalized with a sulfonate group. Trimers such as ETE-S can be added to the CP solution and subsequently attached to the PEDOT-S to increase NW stability. Incorporation can be done by applying an electrical bias (electrofunctionalization) or by applying an iron (III) catalyst (chemical functionalization)^18^. To electrofunctionalize ETE-S onto the PEDOT-S, an electrical bias is applied to the polymer while it is still wet and freshly added to the pores. For contacting, the TE membrane is gently placed onto a conductive tape serving as the anode, and a AgCl dip-in electrode is used as the counter electrode. A bias of 0.9 V was applied between the electrodes throughout the electrofunctionalization. After drying overnight, ethanol was used to remove any conductive tape residues and excess polymer from the surface of the sample after delamination. NWs electrofunctionalized with ETE-S are shown in Figure 2K–M. While we present the incorporation of ETE-S into the NWs, more recent ETE derivatives harboring other chemical moieties (e.g., zwitterionic phosphatidylcholine or positively charged trimethyl ammonium^24^) can also be used to alter NW chemistry using the same methodology.

**Figure 2:**
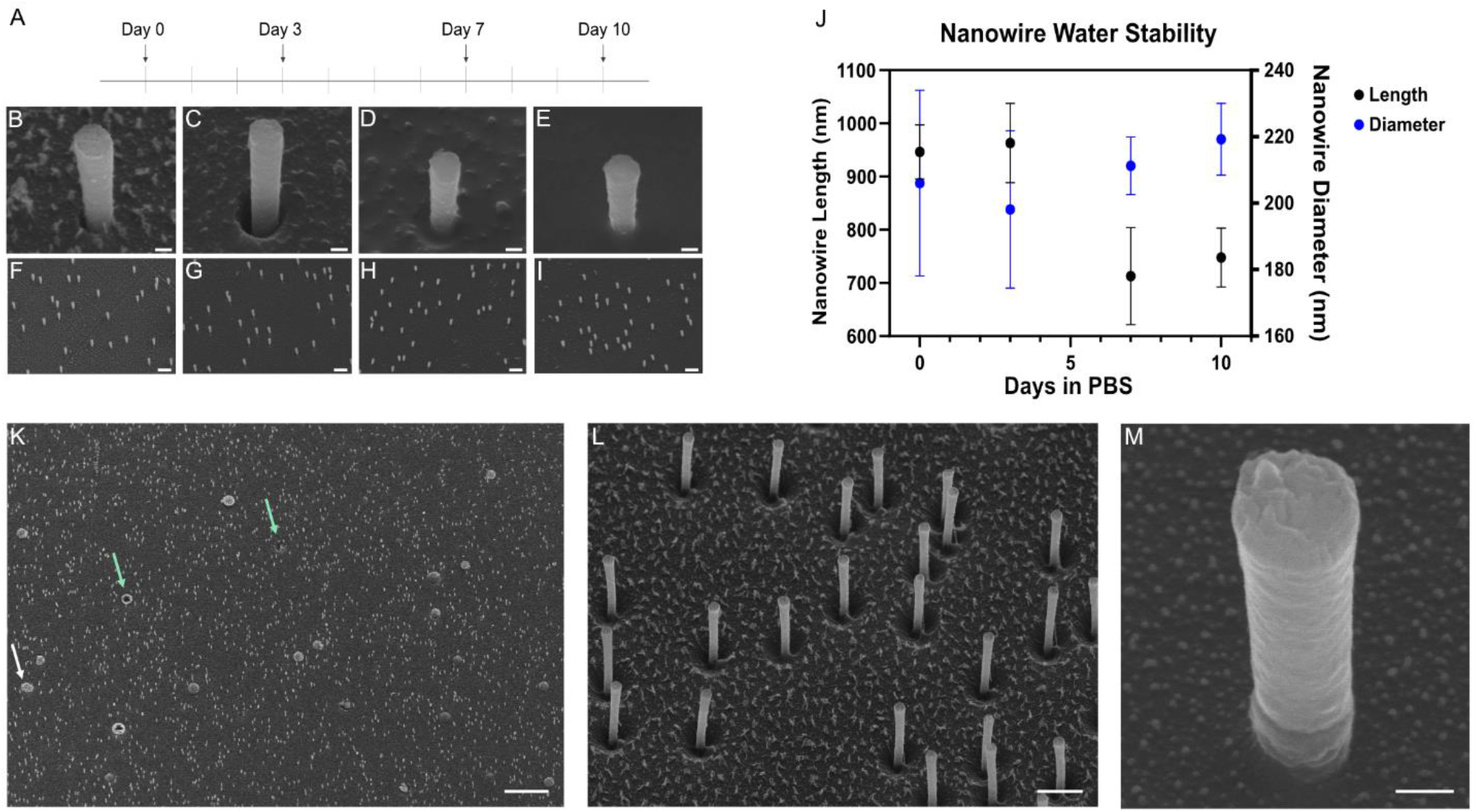
NW stability can be increased by incorporating trimers and electrofunctionalization or chemical strengthening. (A–J) NW stability after being submerged in PBS up to 10 days. (A) Timeline of PBS exposure to NWs. (B–E) 30° tilted SEM images of NW strengthened with trimers and 100 mM Fe^3+^. NWs were imaged before exposure to PBS (B, F), after three days of exposure (C, G), after 7 days of exposure (D, H) and after 10 days of exposure (E, I) Lower magnification images of NWs after exposure available in Figure S6. (J) Graph illustrating NW diameter and NW in length at various time points. Dots on the graph represent the mean, while error bars represent the standard deviation. (K–M) 30° tilted view SEM images of NWs electrofunctionalized with ETE-S. In K, the white arrow indicates the plasticizers inherent to the TE membrane, and the green arrows indicate the holes left from the plasticizers. The scale bars are 100 nm for panels B–E and M, 1 µm for panels F–I and L, and 10 µm for panel K.

While the addition of cations (such as Fe^3+^) or electrofunctionalization with trimers can be used to increase stability, maximum stability was achieved by using a combination of both cations and trimer incorporation (schematic in Figure S5). Since the higher ion concentration increases the viscosity of the polymer solution, iron ions were added after the initial pore filling with PEDOT-S mixed with ETE-S. This allowed the diffusion of iron ions into the pores to simultaneously cross-link the polymer and aid in trimer functionalization. Images of such NWs at day 0 and at different time points after exposure to PBS are shown in Figure 2B–I. For the first 3 days, no decrease in NW diameter or length was observed. After 7 days, the length decreased by 25 % (946 nm day 0 vs 713 nm day 7, p-value < 0.0001), while the diameter did not significantly decrease, Figure 2J. This change in height could be due to NW decomposition, but there is also the possibility that height is decreased due to the accumulation of salt from the PBS on the TE membrane. Summarizing, these experiments have shown that the PEDOT-S NWs can be made water stable by adding cations and trimers.

### Electrical characterization

The NWs remain conductive after processing as mapped by conductive atomic force microscopy (cAFM). NW studied were composed of 5 mg/ml PEDOT-S, and 5 mM Fe^3+^ in PBS. In cAFM, a Pt/Ir coated tip is scanned across the sample, resulting in the simultaneous acquisition of topographic information and electrical current flow between the tip and the biased sample.

Samples with either NWs lying flat on a surface (Figure 3A–D) or NWs extending from a PI membrane (Figure 3E–H) were mapped. For the NWs lying down, NWs were stochastically placed onto pre-patterned Au lines on glass. The Au lines were contacted, allowing an electrical current to flow between the tip and the NWs. Distance dependent conductivity was found, with sections of the NW closer to the gold electrode showing a higher current, Figure 3D.

**Figure 3:**
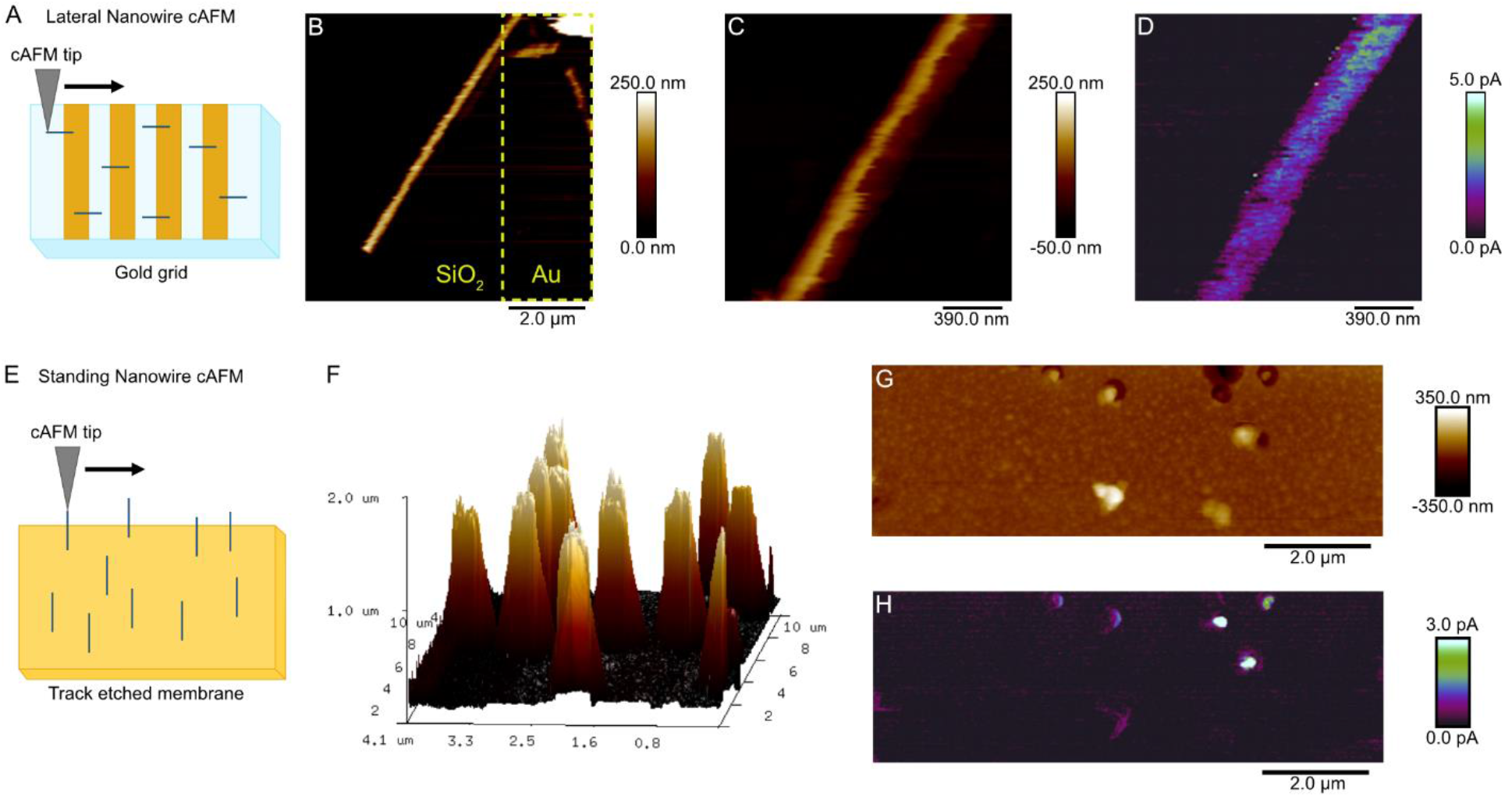
Physical characterization of NWs by conductive atomic force microscopy (cAFM). Panels A and E are diagrams illustrating cAFM configuration, panels B–D are measurements taken on NWs lying flat on a surface, while panels F–H are measurements on standing NWs still embedded in a TE membrane. Topography images of height, both in 2D (B, C, G) and 3D (F) representation, range from white at the highest point, brown at midpoint, to black at the lowest point. Current maps (D, H) express light green at the most conductive point, then ranges through blue, and purple at the midpoints, then to black at the least conductive point. (B) Topography image of NW height. The image shows the NW lying on a patterned surface, part of the NW on the gold section, and the rest of the NW on the glass surface. Gold section is denoted by the dotted yellow line, while the rest of image is glass surface. (C–D) Topography image of height (C) and current map (D) on a section of the same NW as shown in B. (F) Three-dimensional representation of cAFM height measurements on NWs. (F, G) Two-dimensional topography image illustrates height (F) and current map illustrates current (G) measurements.

For NWs standing up (Figure 3E), a thin layer of silver paint was used to contact the bottom of the membrane. The cAFM probe was then used to scan on top of the standing spikes. Scanning on standing wires poses a significant challenge for cAFM due to the high aspect ratio, which can lead to bending or breakage of the NWs when they contact the probe tip. For this reason, we used shorter NWs. Currents in the low pA regime were detected when scanning over the NWs, Figures 3D, S7. In comparison, no current over noise level was registered when the tip was on the naked polymer membrane.

In the lateral configuration, we observed current flow multiple µm away from the contact, Figure 3D. In the vertical configuration, the current flows from the tip through the NW inside the pores before reaching the back contact, thereby extending 12 µm in total. Importantly, this highlights that the polymer inside the pores is electrically continuous throughout its length. Compensating for the large distance, a higher voltage was used (10 V) in the vertical configuration as opposed to the lateral one (5 V). Further optimization of contact chemistry and geometry will likely decrease the observed resistance since PEDOT-S in two-dimensional transistor setups have shown high conductivity^26, 17^. Taken together, the cAFM results show that the NWs are conductive and can be contacted on the backside of the membrane.

### Cellular Interfacing

After establishing the possibility to generate water-stable and conductive NWs, we interfaced them with living cells. The NW membrane was fixed to the bottom of a biocompatible polycarbonate tube, forming an easy-to-handle cell culture vessel. Two vastly different cell types were explored in separate experiments: the single cell algal model system *Chlamydomonas Reinhardtii* and clinically relevant human primary hematopoietic stem and progenitor cells (HSPCs, CD34+), Figures 4 and S8. To enable high-resolution SEM imaging of the NW–cell interface, samples were carefully fixed, dehydrated, and point-dried to retain their native morphology and minimize the risk of geometrical changes.

**Figure 4:**
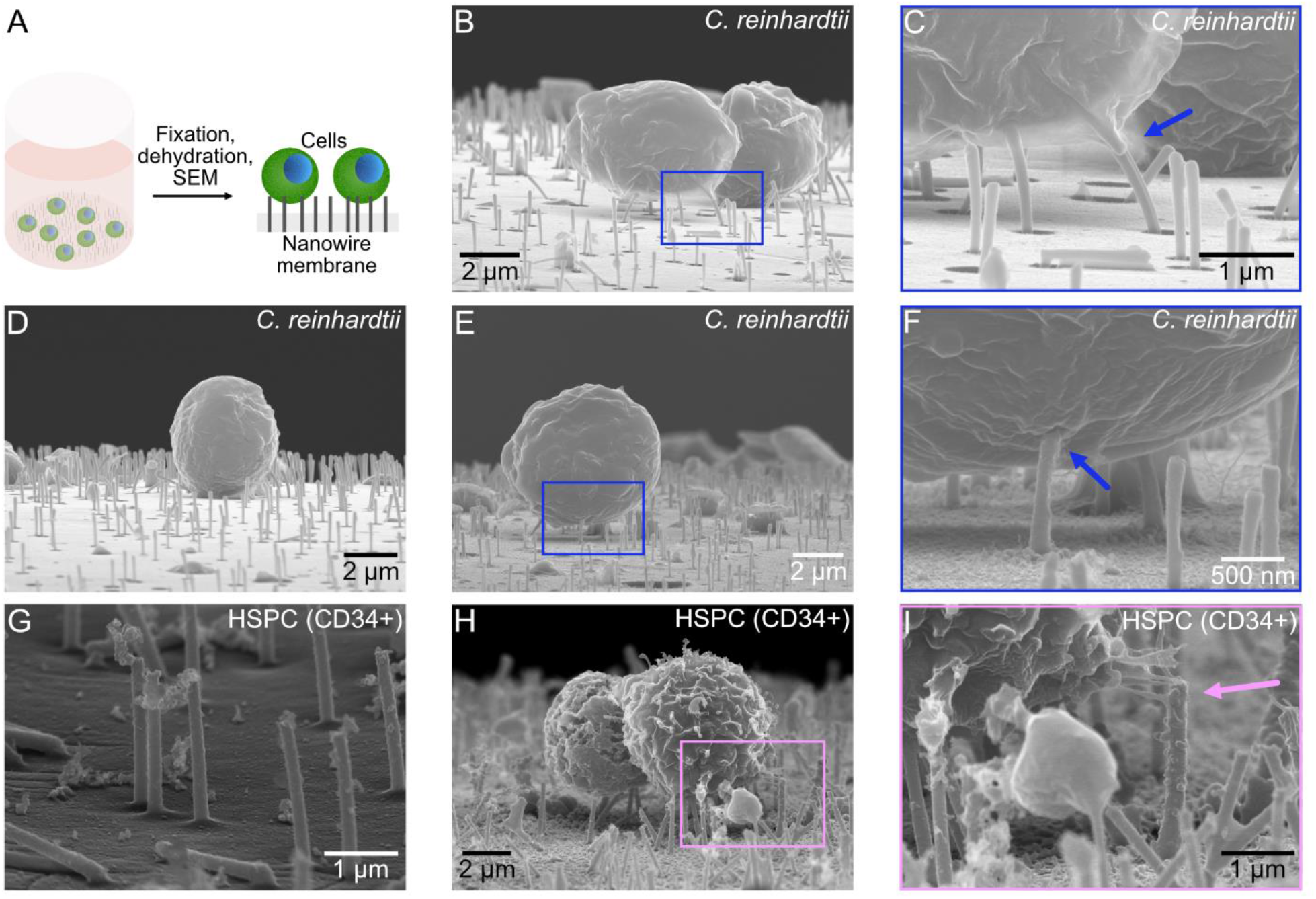
Interfacing living cells with CP NWs stabilized by 10 mM Fe^3+^. (A) schematic depicting cells added to a culture vessel incorporating the NW membrane in the bottom. Following fixation and dehydration, the sample is imaged from the side to reveal the NWs interfacing with the cells. (B–F) SEM images showing algal cells, *C. reinhardtii*, on the NWs. Panels C, F, I are zoomed in from the marked regions in B, E, H. Blue arrows show NWs seemingly reaching inside the cells. (G–I) SEM images showing human primary hematopoietic stem cells (HSPCs, CD34+) interfacing NWs. Image G was obtained in a cell-free region. The pink arrow in I depicts cells reaching out to grab the spikes.

Interfacing nanowires with the sweet water algae *C. reinhardtii* poses a challenge for the NWs’ water stability since the cells’ preferred medium is hypo-osmolar and will more readily degrade the NWs. The cells were dispersed in dilute PBS (20 %, in water), added to the culture wells, and centrifuged onto the NWs at 600 g to achieve a controlled pelleting without risk of cell aggregation. Despite the centrifugation induced pressure, SEM images revealed NWs extending vertically in areas with and without cells, Figure 4B–F. Some broken NWs were observed, which could be related to mechanical impact from the cells, fluid exchanges during fixation, or dehydration. Interestingly, NWs close to the cells bent towards the cell membrane and were seemingly reaching intracellularly, Figure 4C, F. Nanostructures of similar dimensions can spontaneously pierce the cell membrane ^22^ and owing to the electrically conductive nature of the NWs, the intimate cell–NW contact can relay external voltage pulses for local destabilization of the cell membrane^27^ thereby enhancing the possibility for intracellular access.

Human primary HSPCs are defined by a surface marker, CD34+, and are routinely used in the clinic as bone marrow transplants. We have previously described how tubular nanostructures, nanostraws, could be used to access the intracellular space of HSPCs without being noticed by the cellular defense systems^6^. Apart from this, little is known about how these cells interface with axially elongated nanostructures despite their clinical relevance. Similar to the experiments with the algae, cells were centrifuged onto the NWs and subsequently prepared for SEM, Figure 4G–I. Importantly, these sensitive cells were dispersed in an isotonic medium. Even after rinsing, fixation, dehydration, and critical point drying, the NWs were still protruding out from the NW membrane. Compared to the algal samples, more debris was observed, possibly being cell residues from the freeze-thaw cycle before being applied to the NWs, or coming from the cell medium. Cell morphology did not show any obvious signs of cell damage. It was repeatedly observed that cells close to NWs extended towards them, seemingly wanting to grab on, Figure 4I, as observed previously for other NW structures^28, 29^. In summary, the CP NWs can be made water stable and can be used to interact with living cells, both human and algal, and form close cell–NW interfaces.

## Conclusions

In summary, we have demonstrated a lithography-free platform to generate PEDOT-S nanowires with controlled dimensions. By avoiding lithography, NWs can be produced at a scale where the ICP reactor governs the sample size (typically a 4’’ or 6’’ sample size, depending on the reactor used). NWs were made water stable through introduction of thiophene trimers and either cations or electrofunctionalization. Even when interfacing the cells with the NWs, they remained intact and could withstand 600 g of cell pelleting.

The NW processing described here is not limited to the use within cellular interfacing but is general in nature. The geometry and high surface-to-volume ratio could allow for making organic electrochemical transistors (OECTs) with very high switching speeds owing to an optimal gating geometry and short diffusion paths for ions moving in or out of the NWs. These qualities open new doors for electrical and biological devices based on one-dimensional CPs.

## Methods

### NW formation

A commercially available poly(3,4-ethylenedioxythiophene)butoxy-1-sulfonate (PEDOT-S) solution (Clevios KS LVW 2012, Hereus Epurio Innovation) was applied onto track-etched (TE) polyimide (PI) and polycarbonate (PC) membranes. TE membranes with different pore geometries were evaluated: diameters ranging from 160–200 nm, pore densities 2– 5.5 e^7^ cm^−2^, and thickness of 12–25 um (It4IP S.A., Louvain-la-Neuve, Belgium).

In some experiments, different salts were added to the polymer to increase the ionic strength of the solution. To ensure a well-dispersed solution, the PEDOT-S was ultrasonicated before adding it to the TE membrane in order to cover both the top of the membrane and to fill up the nanopores. The polymer solution was left to dry overnight followed by mechanical removal of excess (dry) polymer residing on top of the TE membrane using masking tape (Heavy Duty Masking tape, 3M). The polymer filled TE membrane was electrostatically or mechanically adhered to a 4’’ Si wafer using an electrostatic gun (Sigma-Aldrich, Zerostat anti-static instrument) or Kapton tape for subsequent semiconductor processing.

### ICP-RIE etching

Oxygen based inductively couple plasma reactive ion etching (ICP-RIE) was used to selectively remove TE membrane in an anisotropic manner (Plasma-Therm, APEX SLR). Processing parameters for standard spikes using ICP-RIE: 50 sccm O2, 25 W RF, 500 W ICP, 10 mTorr reactor pressure, 120-180 s etch duration. Processing parameters for a ‘strong’ etch using ICP-RIE: 50 sccm O2, 50 W RF, 600 W ICP, 10 mTorr reactor pressure, 180 s etch duration. The duration of this etching step governs the length of the finished NW and was varied to extract an etch rate.

### Polymer solution for electrofunctionalization and chemical functionalization

Conductive polymer solution was prepared by dissolving 10 mg/ml 4-(2-(2,5-bis(2,3-dihydrothieno[3,4-b][1,4]dioxin-5-yl)thiophene-3-yl)ethoxy)butane-1-sulfonate (ETE-S), 5 mg/ml PEDOT-S (PEDOT-S; Sample: Clevios KS LVW 2012, Hereus Epurio Innovation), and iron (III) sulfate hydrate (Sigma-Aldrich; Final concentration in solution 5 mM) in phosphate buffered saline (PBS). For chemical functionalization, CP solution only contained 10 mg/ml ETE-S, 5 mg/mL PEDOT-S in PBS. To ensure a well-dispersed and low viscosity solution, samples were ultrasonicated 5 times (20 pulses at 100 % amplitude and 0.75 cycle; Hielscher).

### Electrofunctionalization

Electrofunctionalization was performed by pipetting CP solution onto a polyimide membrane which was adhered to a piece of conductive copper tape. A Keithley Sourcemeter (Model 2612B) was used to apply a 0.9 V electrical bias in between the CP solution (AgCl dip-in electrode) and the copper tape for 30–60 minutes. The resulting current was monitored throughout the electrofunctionalization process. After electrofunctionalization sample was allowed to air dry overnight.

To remove the copper tape, the membrane was submerged in 99.5 % ethanol (Solveco). Possible adhesive residues were removed from the back of the membrane by using a Kimwipe (Kimtech) and 99.5 % ethanol. Excess alcohol was left to evaporate. Excess CP was mechanically delaminated using tape (Heavy Duty Masking tape, 3M). The top surface was cleaned using a Kimwipe with 99.5 % ethanol to remove possible contaminants. Following this, the electrofunctionalized samples were processed using standard ICP-RIE as described before.

### Chemical functionalization

CP solution was pipetted onto a PI membrane and allowed to dry for an hour, before the addition of 100 mM iron (III) sulfate hydrate (Sigma-Aldrich). Sample was allowed to dry and further processed as a standard sample.

### SEM Imaging

For scanning electron microscopy (SEM), small pieces from the NW membrane were cut out and sputter coated (Q150T ES, Quorum) with 5 nm of Pt:Pd (80:20) alloy. Samples were imaged using cold-field emission SEM (SU8010, Hitachi) operating at 10 kV. NW diameter and NW length were measured in ImageJ software (Version 1.54g) using a 30° tilted view 10,000x magnification SEM image of each sample. Measured length was multiplied by a factor of two, to accommodate for tilt. N for each sample point varies between 14 and 33, all NWs completely visible, intact, and in frame of the image were measured. For SEM of cells and algae on NWs, additional sample preparation was needsed.

Human primary stem and progenitor cells (CD34+) cells were dispersed in PBS or algal cells from the strain *Chlamydomonas Reinhardtii* dispersed in dilute PBS (0.2x) were centrifuged onto the NW membrane followed by fixation in 2 % glutaraldehyde, 2 % formaldehyde in PBS. After fixation, cells were stored in Sorensen’s phosphate buffer. An ethanol exchange series of 10 min each in 30 %, 50 %, 75 %, 90 %, 95 %, and 99.5 % ethanol was used. Critical point drying (CPD) was used to completely remove the ethanol from the cells with minimal disruption of the cellular shape in a Quorum K850 CPD (Quorum Technologies). The CP dried samples were coated with 5 nm Pt:Pd and imaged using SEM.

### NW stability in phosphate buffered saline

Samples were taped into a Petri dish (Avantor), which was then filled with 3 ml of PBS. This formed a layer of PBS over the samples. PBS was removed from Petri dish, by gently tilting and pouring out the PBS. In order to remove residual PBS, 5 rinses of 1 ml milli-Q water was done, then 3 rinses of 1 ml of 99.7 % ethanol was done. Excess liquid from each rinse was removed by tilting the Petri dish, and using a Kimwipe to absorb liquid at the edge of the dish if the solution does not readily evaporate. Ethanol was allowed to evaporate after the last rinse. Visualization of NWs was done via SEM as described in the ‘SEM imaging’ section.

### Conductive Atomic Force Microscopy

C-AFM was performed on a Dimension Icon XR equipped with a PF-TUNA module (current sensitivity 20 pA/V) from Bruker. Pt/Ir-coated silicon probes (k = 3 N/m) were used to simultaneously map topography and current in PF-TUNA mode. The current maps were obtained by constantly biasing the Au substrate, while keeping the scanning AFM probe at ground. All the measurements were performed at room temperature in ambient atmosphere.

### Cell culture

Stem and progenitor cells: Umbilical cord blood was collected at Skåne University Hospital and Helsingborg Hospital. Mononuclear cells were extracted using Lymphoprep tubes (Alere Technologies, no. 1019818). CD34+ cells from mixed donors were isolated from mononuclear cells with a CD34 MicroBead Kit (Miltenyi Biotec, no. 130–046-703) according to the manufacturer’s instructions. Cells were cells were frozen in 90 % fetal calf serum + 10 % dimethyl sulfoxide before use, and then thawed for the experiment. After thawing, cells were cultured using StemSpan serum-free expansion medium (Stemcell Technologies, no. 09650) with human stem cell factor (PeproTech, no. 300–07), human thrombopoietin (PeproTech, no. 300– 18), and human recombinant Human Flt3-Ligand (PeproTech, no. 300–19), 100 ng/ml, 37 °C, 5% CO2. Cells were washed in PBS and centrifuged at 150 g for 10 min to remove debris. The cell pellet was resuspended in PBS and centrifuged onto NWs at 600 g for 3 min.

Algal Cells: A cell suspension of the model cell line *Chlamydomonas reinhardtii* was suspended in 20 % PBS and centrifuged at 350 g for 10 min to remove debris. The cell pellet was resuspended in 20 % PBS and centrifuged onto NWs at 600 g for 3 min.

## Supporting information

File containing Supplemental Figures

## Ethics Statement

Work with primary human CD34+ cells was approved by the regional ethical committee for Lund/Malmö (Regionala Etikprövningsnämnden i Lund/Malmö), approval no. 2010–696. Informed consent was obtained from mothers of the umbilical cord blood donors, and all samples were deidentified prior to use in the study.

## Acknowledgments

This study was accomplished within the Lund University Strategic Research Areas MultiPark and NanoLund. We thank Dr. Maria Svensson Coelho (Lund University) for donating *C*.

Reinhardtii cells, and Armin Dardashti for experimental assistance. Equipment within Lund Nano Lab (LNL) was used to enable this research.

## Author contributions

Conceptualization: MH Monomer synthesis: AHM, MAS Funding acquisition: RO, MH

Experiments and reagents: DH, UA, LS, FE, CM, MB, ML Cell culturing: LS, JL

Supervision: MH Writing–original draft: DH, MH

Writing–review & editing: All authors reviewed and edited the draft

## Data and materials availability

All data are available in the main text or the supplementary materials.

## Supplementary Materials

Supplementary Figures S1–S8

