## Supplementary material for "Lithography-free Water Stable Conductive Polymer Nanowires": File containing Supplemental Figures

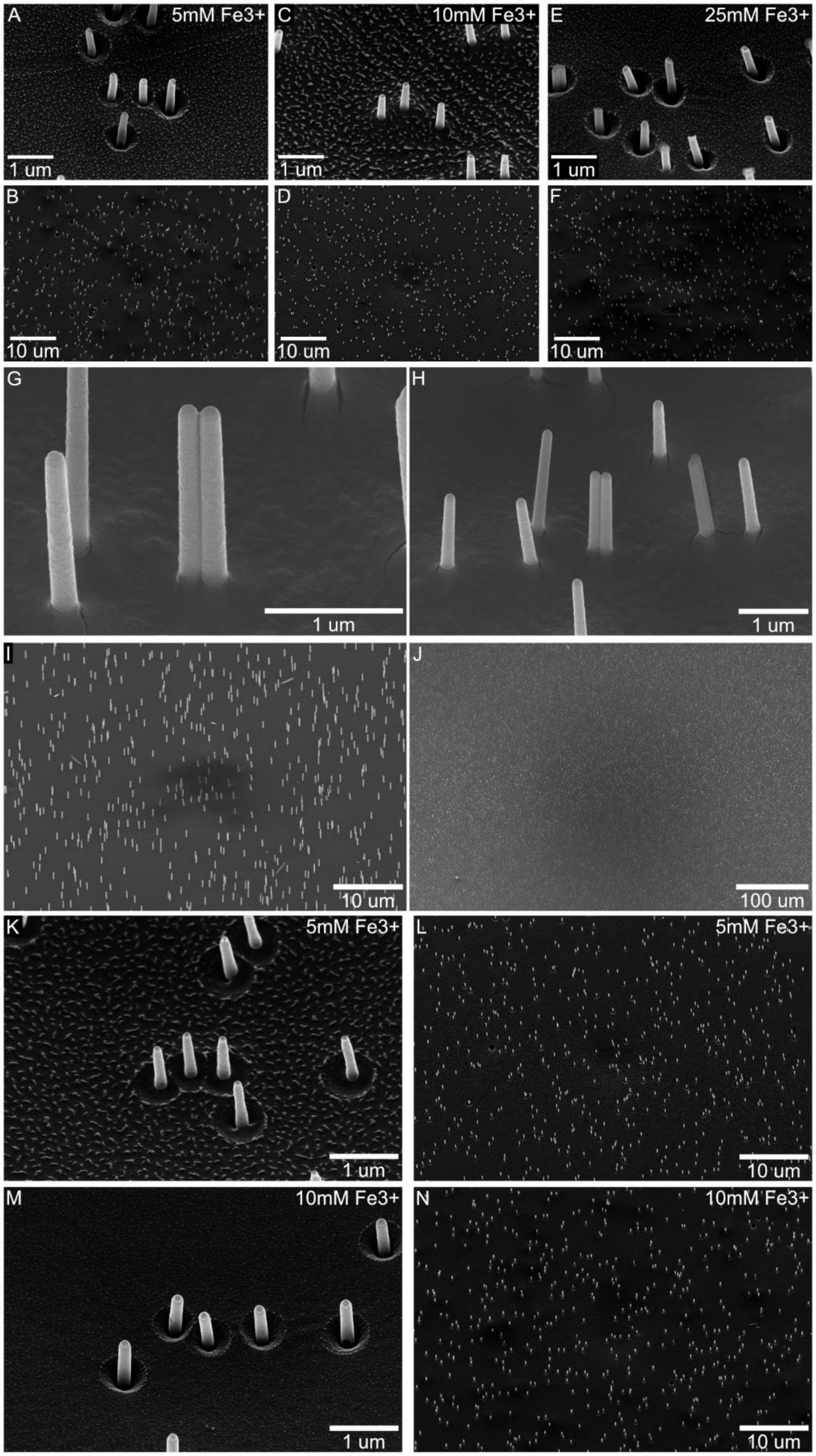


Figure S1: 30° tilted view SEM images depicting nanowires made with varying amounts of $Fe^{3+}$ added during templating. PEDOT-S concentration used was 5 mg/ml for A–J, and 10 mg/ml for

K–N. A, B are made with 5 mM $Fe^{3+}$; C, D are made with 10 mM $Fe^{3+}$; E, F are made with 25 mM $Fe^{3+}$. Panels G–J include 30° tilted view SEM images depicting nanowires going from high to low magnification. Please note the excellent uniformity over large areas. Nanowires are made with 5 mg/ml PEDOT-S and 20 mM $Fe^{3+}$. Panels K–N include 30° tilted view SEM images with nanowires made with 10 mg/ml PEDOT-S with 5 mM $Fe^{3+}$ (K, L), and with 10 mg/ml PEDOT-S with 10 mM $Fe^{3+}$ (M, N).

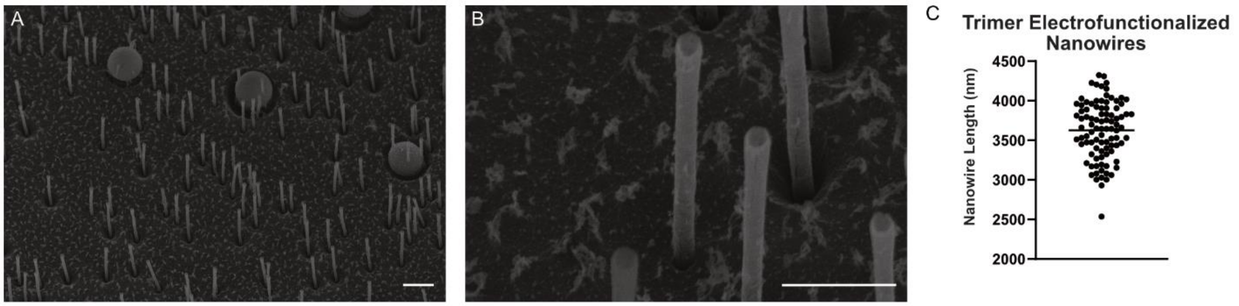


Figure S2: 30° tilted view SEM images of nanowires grown up to 4 μm using trimer incorporation and electrofunctionalization. Nanowires did not undergo ethanol rinse after delamination. (A) Scale bar 2 μm in panel A, and 1 μm in panel B. (C) Graph of observed lengths of nanowires in panel A. Variation in nanowire observed length is likely due to deviations in viewing angle. Each dot represents one nanowire, n = 93. Line illustrates mean.

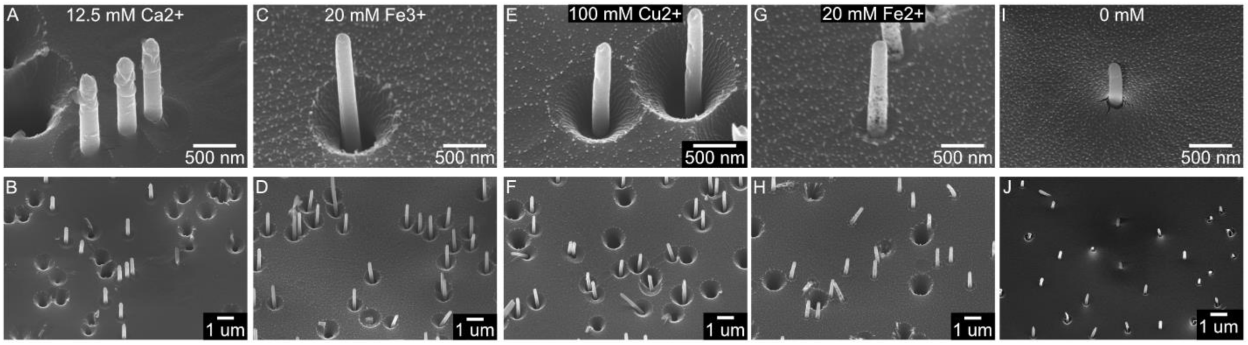


Figure S3. 30° tilted view SEM images depicting nanowires made with different ions added during templating (non-optimized settings). Nanowires in panels A, B are made with 12.5 mM $Ca^{2+}$; C, D are made with 20 mM $Fe^{3+}$; E, F are made with 100 mM $Cu^{2+}$; G, H are made with 20 mM $Fe^{2+}$; I, J are made without any additions.

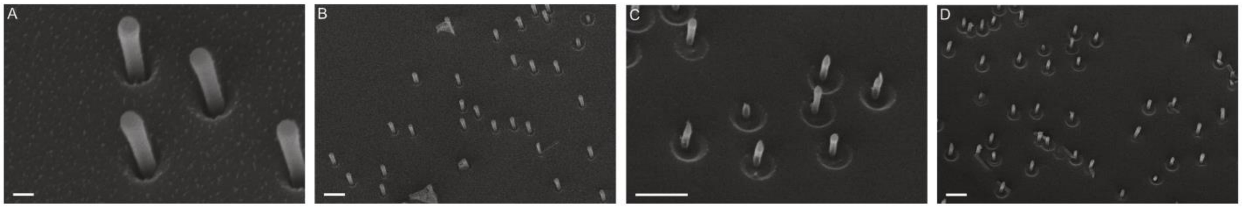


Figure S4: 30° tilted view SEM images illustrating PBS stability of nanowires composed of PEDOT-S with low levels of iron. Nanowires were composed of 5 mg/ml PEDOT-S and 5 mM $Fe^{3+}$. SEM images depicting nanowires before PBS exposure (A–B) and exposed to PBS for 24 hours (C–D). In panels C, D, degradation of nanowire tips is visible, as well as multiple displaced nanowires. For panel A Scale bar is 200 nm, for panels B–D scale bars are 1 µm.

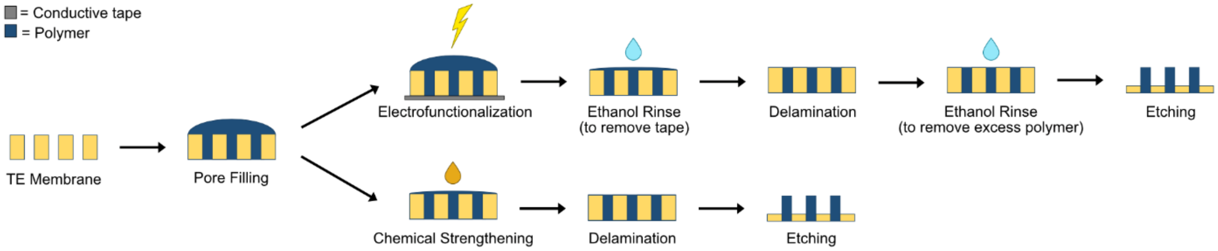


Figure S5: Schematic side-view describing incorporation thiophene trimers into nanowires and subsequent processing. A track-etched (TE) membrane is filled with conductive polymer solution. The sample then either undergoes electrofunctionalization or chemical strengthening. For electrofunctionalization, a 0.9 V bias is applied via piece of conductive tape, with a AgCl dip-in electrode as the counter electrode for 30-60 min. Sample is then allowed to dry overnight. An ethanol rinse is used to remove the sample from the conductive tape, and remove residual adhesive. After ethanol has evaporated, the sample is then delaminated to remove the top layer of polymer. Sample is then rinsed again with ethanol, to remove residual polymer. The TE membrane is then etched by oxygen via inductively coupled plasma reaction ion etching (ICP-RIE) to reveal the nanowires. For chemical strengthening, a solution of 100 mM $Fe^{3+}$ is added to the surface of the sample. Sample is then allowed to dry overnight. Sample is then delaminated, then etched via standard ICP-RIE etching. Please note that schematic is not to scale.

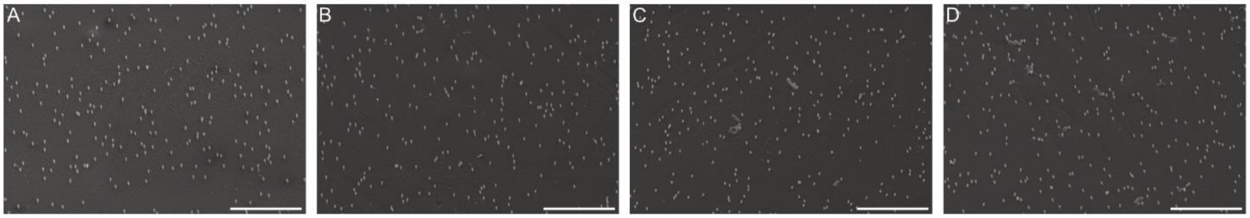

Figure S6: 30° tilted view SEM images of nanowire stability after exposure to PBS. Nanowires are stable in PBS up to 10 days. SEM images are from the same respective samples as Figure 2B–I, only imaged at a lower magnification. (A–D) Nanowires were imaged before exposure to PBS (Day 0, A), after three days of exposure (Day 3, B), after 7 days of exposure (Day 7, C) and after 10 days of exposure (Day 10, D). Scale bars are 10 microns.

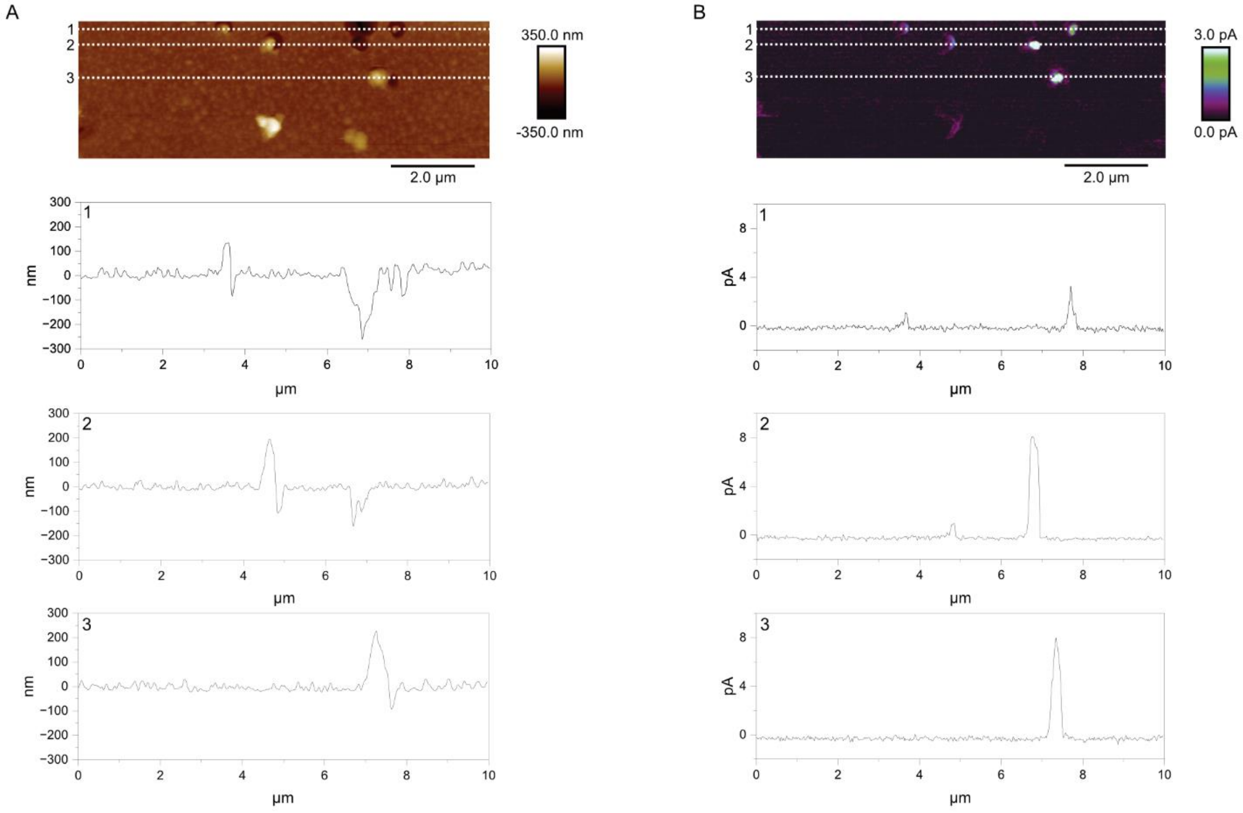


Figure S7: CAFM data for standing nanowires. Panel A is a topography image illustrating height and panel B is current map demonstrating current (B) together with section profiles along the lines marked in the maps (1-3).

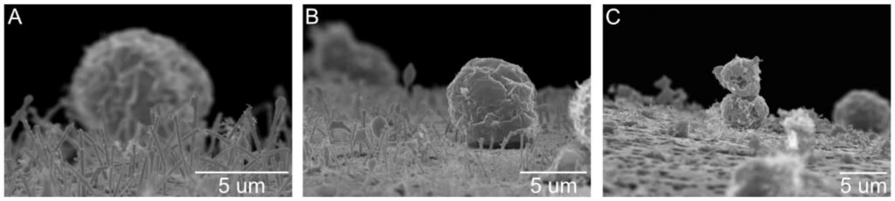

Figure S8. SEM images depicting CD34+ cells on nanowires (stabilized with 10mM $Fe^{3+}$). A, B depicts cells after successful dehydration and critical point drying whereas C shows cells after unsuccessful SEM preparation/dehydration.
